# Direct visualization of single nuclear pore complex proteins using genetically-encoded probes for DNA-PAINT

**DOI:** 10.1101/579961

**Authors:** Thomas Schlichthaerle, Maximilian T. Strauss, Florian Schueder, Alexander Auer, Bianca Nijmeijer, Moritz Kueblbeck, Vilma Jimenez Sabinina, Jervis V. Thevathasan, Jonas Ries, Jan Ellenberg, Ralf Jungmann

## Abstract

The Nuclear Pore Complex (NPC) is one of the largest and most complex protein assemblies in the cell and – among other functions – serves as the gatekeeper of nucleocytoplasmic transport. Unraveling its molecular architecture and functioning has been an active research topic for decades with recent cryogenic electron microscopy and superresolution studies advancing our understanding of the NPC's complex architecture. However, the specific and direct visualization of single copies of NPC proteins and thus the ability to observe single-molecule heterogeneities of these complex structures is thus far elusive. Here, we combine genetically-encoded self-labeling enzymes such as SNAP-tag and HaloTag with DNA-PAINT microscopy. We employ the high localization precision in DNA-PAINT and molecular contrast of these protein tags to optically resolve single copies of nucleoporins in the human Y-complex in three dimensions with a precision of ~3 nm. This technological advancement now enables structural studies of multicomponent complexes on the level of single proteins in cells using optical fluorescence microscopy.

---

Super-resolution techniques allow diffraction-unlimited fluorescence imaging^1,2^ and with recent advancements, true biomolecular resolution with an all-optical microscopy setup is well within reach^3-5^. One implementation of singlemolecule localization microscopy (SMLM) is called DNA points accumulation in nanoscale topography^4^ (DNA-PAINT), where dye-labeled DNA strands (called ‘imager’ strands) transiently bind to their complements (called ‘docking’ strands) on a target of interest, thus creating the typical ‘blinking’ used in SMLM to achieve super-resolution. While localization precisions down to approximately one nanometer (basically the size of a single dye molecule) are now routinely achievable from an imaging-technology perspective, this respectable spatial resolution has yet to make its way into widespread applications in cell biological research. Currently, this is mainly hampered by the lack of small and efficient protein labels. Recent developments of nanobody- or aptamer-based tagging approaches^6-10^ are providing an attractive route ahead, however both approaches are not yet deploying their full potential either due to limited binder availability (in the case of nanobodies) or lack of large-scale analysis of suitable super-resolution probes (in the aptamer case).

While we are convinced that some of these issues might be resolved in the future, we here introduce the combination of widely-used, genetically-encoded self-labeling enzymes such as SNAP-tag^11^ and HaloTag^12^ with DNA-PAINT to enable 1:1 labeling of single proteins in the Nuclear Pore Complex (NPC) using ligand-conjugated DNA-PAINT docking strands. The NPC is responsible for the control of nucleocytoplasmic transport and a highly complex and sophisticated protein assembly. NPCs contain multiple copies of approximately 30 different nuclear pore proteins called nucleoporins (NUPs) and have an estimated total molecular mass of ~120 MDa, placing NPCs among the largest cellular protein complexes^13^. Due to their diverse function in controlling molecular transport between the nucleus and the cytoplasm (a necessity for example for gene expression and cell growth), NPCs are a major target for structural biology research with characterization by e.g. cryogenic electron microscopy^13^ (cryo-EM) or optical super-resolution techniques^14,15^. State-of-the-art cryo-EM studies^16-18^ – reaching impressive pseudo-atomic resolution – have advanced our structural understanding in recent years. It is now possible to not only elucidate how NUPs in NPCs are arranged, but also to shed light on how structural changes of NPCs are connected to their dysfunction^19^. However, even with recent advancements in cryo-EM instrumentation, molecular specificity necessary to resolve single NUPs in NPCs proves still elusive, mainly due to the lack of high protein-specific contrast. As single-particle class averages of thousands of NPCs are usually necessary to obtain high-resolution structures, uncovering molecular heterogeneities of single NUPs in different NPCs is virtually impossible. Fluorescence-based techniques on the other hand offer exquisite molecular contrast and specificity due to the use of dye-labeled affinity reagents targeting single protein copies in cells. However, until recently, the necessary resolution to spatially resolve single small proteins in a larger complex has not been achieved due to limitations in labeling (small and efficient probes) and imaging technology (providing sub-10-nm spatial resolution).

Here, we combine DNA-PAINT microscopy with small, genetically-encoded self-labeling enzymes such as SNAP- and HaloTag to overcome limitations in optical super-resolution microscopy. We present a straightforward protocol to target these tags in a variety of engineered cell lines using the DNA-conjugated ligands benzylguanine (BG) and chloroalkane against SNAP-tag^11^ and HaloTag^12^, respectively (**Fig. 1a** and **b**). We investigate the achievable labeling precision and reduction of linkage error of SNAP-tag and HaloTag compared to traditional primary and DNA-conjugated secondary antibody labeling and furthermore examine their performance in contrast to DNA-conjugated nanobodies against GFP fusion proteins in single NPCs. Finally, using our optimized labeling and imaging protocol, we optically resolve – for the first time – single copies of NUP96 proteins in the Y-complex of the NPC, spaced only ~12 nm apart using optical super-resolution microscopy. This now moves super-resolution microscopy into a sweet spot to investigate questions in structural biology with unprecedented spatial resolution and molecular specificity for a large variety of target molecules in genetically-engineered cell lines.

**Figure 1.**
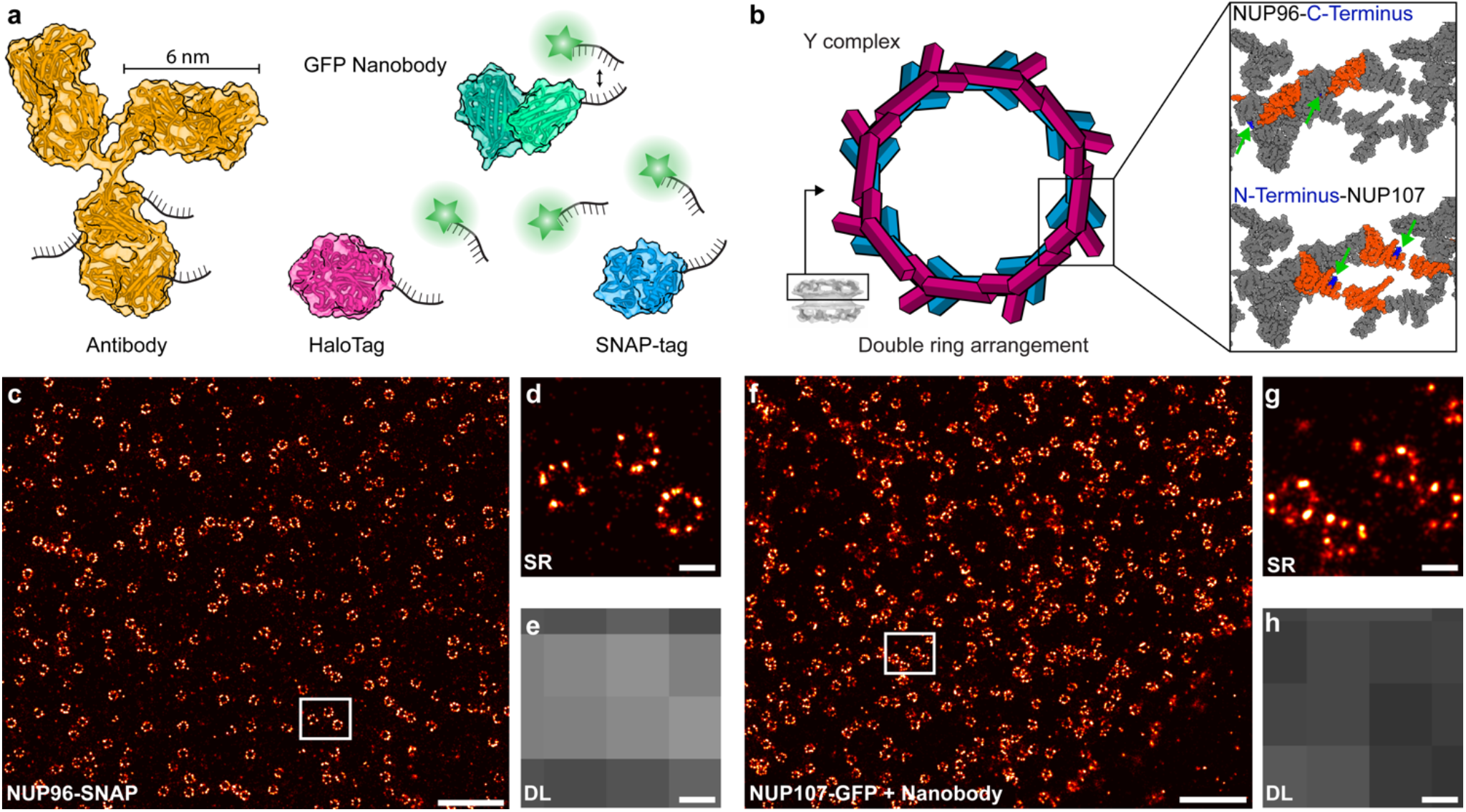
DNA-PAINT labeling probes to target NUPs in NPCs. (**a**) Comparison of different labeling probes (Secondary Antibody: Yellow, GFP Nanobody: green, HaloTag: magenta, SNAP-tag: blue) conjugated with DNA strands for DNA-PAINT imaging (Cartoons are based on Protein Database (PDB) entries: Secondary Antibody (1IGT), GFP Nanobody (3K1K), HaloTag (4KAF), SNAP-tag (3KZZ)). Proteins are to scale. (**b**) NPCs contain 16 copies of NUP96 and NUP107 in the cytoplasmic as well as the nuclear ring. **Top right:** C-terminally-labeled (blue, highlighted by green arrows) NUP96 structure (orange) highlighted in the zoom-in of a symmetry center on the ring (~12 nm apart). **Bottom right:** N-terminally-labeled (blue, highlighted by green arrows) NUP107 structure (orange) in the zoom-in of a symmetry center on the ring (~12 nm apart). Distances and cartoons derived from PDB entry: Nup(5A9Q). (**c**) DNA-PAINT overview image of NUP96-SNAP in U2OS cells. (**d**) Super-resolved zoom-in of highlighted area in (**c**) reveals the arrangement of NUP96 in NPCs. (**e**) Diffraction-limited image of the same area. (**f**) DNA-PAINT overview image of NUP107-GFP in HeLa cells. (**g**) Super-resolved zoom-in of highlighted area in (**f**). (**h**) Diffraction-limited image of the same area. Scale bars: 5 μm (**c, f**), 100 nm (**d, e, g, h**).

To implement genetically-encoded self-labeling tags for DNA-PAINT microscopy, we first assayed our ability to use BG-modified docking strands to target SNAP-tags C-terminally fused to NUP96 proteins in U2OS cell lines created by CRISPR/Cas9 engineering^20^. Labeling was performed post-fixation and -permeabilization using standard labeling protocols^15^ adapted for DNA-PAINT (see **Online Methods**). The resulting 2D DNA-PAINT super-resolution image is shown in **Figure 1c**. A zoom-in and comparison to the respective diffraction-limited image reveals the expected 8-fold symmetry of NUP96 proteins in the super-resolution micrograph (**Fig. 1d, e**). We then performed labeling of NUP107-GFP fusion proteins using a DNA-conjugated anti-GFP nanobody^21^ and obtained qualitatively similar results (**Fig. 1f-h**).

To evaluate labeling quality and precision in a quantitative manner, we next compared results of more traditional labeling of NUP96 using primary-secondary antibodies to those of NUP96-SNAP, NUP96-Halo, NUP107-SNAP, and NUP107-GFP cell lines targeted with their respective ligands. The NPC architecture presents a well-suited model to benchmark novel labeling approaches with regards to overall labeling efficiency and limits of spatial resolution, in a sense similar to an *in vitro* DNA origami calibration standard^22^, but inside a cell. Previous EM studies revealed that NUP96 and NUP107 proteins are present in the Y-complex, which forms the cytoplasmic as well as nuclear NPC double ring arrangement in an 8-fold symmetry. The two double rings are spaced ~50 nm apart with each side containing 16 protein copies^16,23^. The two copies of the proteins in each symmetry center are arranged in Y-complexes spaced ~12 nm apart (**Fig. 1b**). In order to quantitatively compare different labeling approaches, we first acquired 2D DNA-PAINT data using identical image acquisition parameters (see **Supplementary Tables 1-3** for details). Next, we selected single NPC structures in the reconstructed super-resolution image, aligned them on top of each other (i.e. the center of the NPC rings, thus creating a sum image) and performed a radial distance measurement over all localizations. This analysis yields two observables for quantitative comparison: First, the average fitted ring radius and second, the width of this distribution. Dissimilar fitted radii for the same protein labeled using different tags are a measure for potential systematic biases introduced by a preferential orientation of the labeling probes. The width of the distribution on the other hand is a proxy for label-size-induced linkage error, i.e. broader distributions originate from ‘larger’ labeling probes. Our data in **Figure 2** provides a quantitative evaluation and comparison of NUP107-SNAP, NUP107-GFP, NUP96-SNAP and NUP96-Halo cell lines targeted with their respective DNA-conjugated labeling probes (see also **Supplementary Figures 1-5**). We furthermore compare our results with NUP96 labeled using primary and DNA-conjugated secondary antibodies in wildtype U2OS cells. We obtained radii of 53.7 ± 13.1 nm for NUP107-SNAP (**Fig. 2a** and **Supplementary Figure 1**) and 54.6 ± 11.9 nm for NUP107-GFP (**Fig. 2b** and **Supplementary Figure 2** and **6b**), as well as 55.9 ± 12.6 nm for NUP96-SNAP (**Fig. 2c** and **Supplementary Figure 3**) and 56.2 ± 10.2 nm for NUP96-Halo (**Fig. 2d** and **Supplementary Figure 4**), in close overall agreement to earlier EM- and fluorescence-based studies^15,16^. Interestingly, for the antibody-stained sample against NUP96 (**Fig. 2e** and **Supplementary Figure 5** and **Supplementary Figure 6a**), we obtained a considerably smaller radius of 43.5 nm. This could be explained by primary and DNA-conjugated secondary antibodies potentially binding preferentially towards the inside of the NPCs. However, not only did the antibody-stained samples yield a smaller apparent NPC radius, also the measured width of the distribution (16 nm) was larger compared to the genetically-encoded tags due to the increased size of the antibodies. In the case of genetically-encoded tags, the width of the distributions is considerably smaller (see also **Supplementary Table 4**) due to the reduced linkage error to the actual protein location, which is in accordance to previous studies which report the introduction of artefacts due to the labeling probes^8,10,24^.

**Figure 2.**
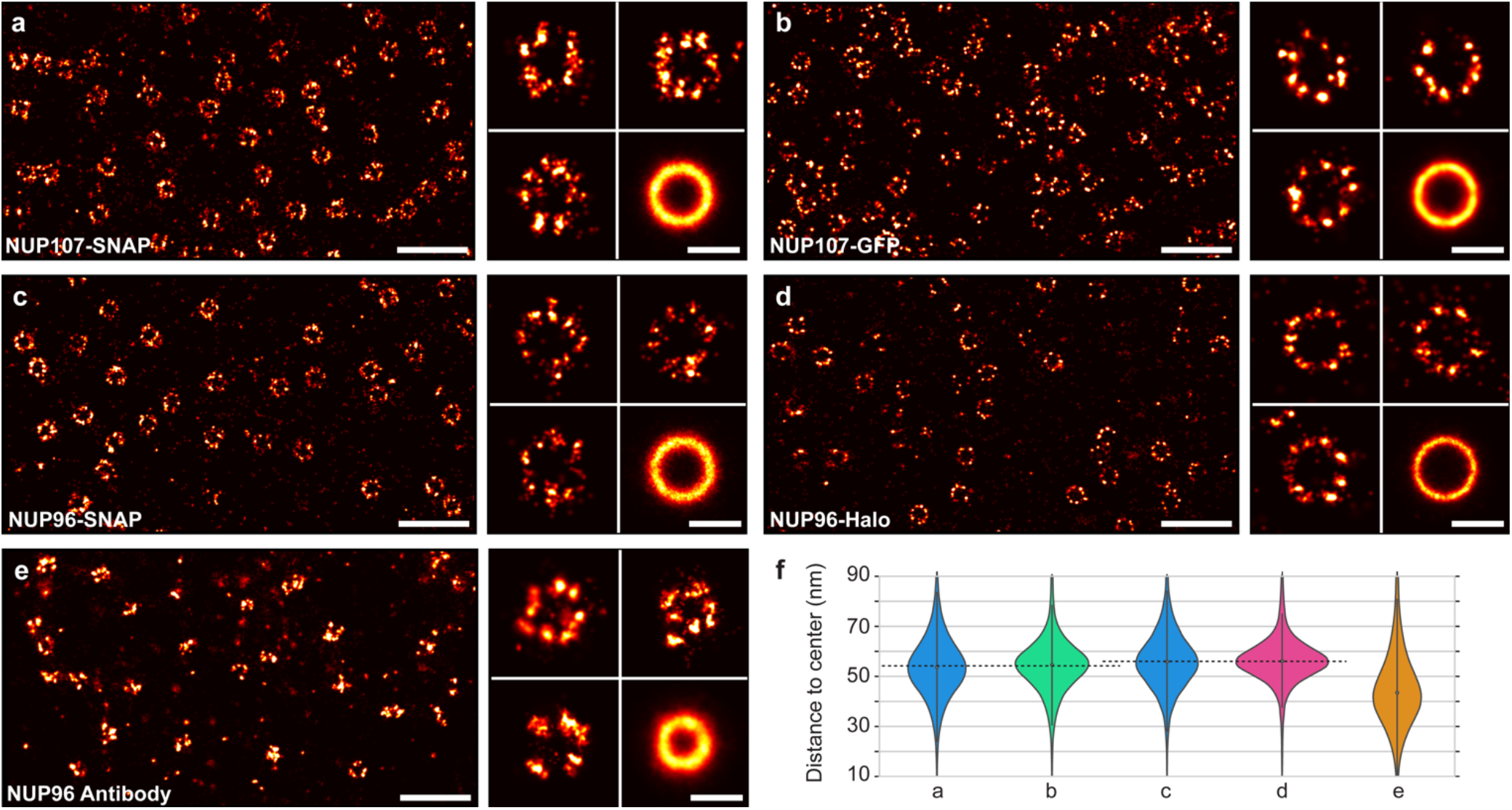
Quantitative comparison of different labels for NUPs. (**a**) NUP107-SNAP overview image (**left**). Zoom-in to individual NPCs and sum image (n = 398) (**right**). (**b**) NUP107-GFP overview image (**left**). Zoom-in to individual NPCs and sum image (n = 486) (**right**). (**c**) NUP96-SNAP overview image (**left**). Zoom-in to individual NPCs and sum image (n = 288) (**right**). (**d**) NUP96-Halo overview image (**left**). Zoom-in to individual NPCs and sum image (n = 191) (**right**). (**e**) NUP96 Antibody overview image (**left**). Zoom-in to individual NPCs and sum image (n = 185) (**right**). (**f**) Violin plots of the distance to the center of localizations. Median radii and standard deviation were obtained for each label: NUP107-SNAP (from **a**, 127773 fitted localizations, 53.7 ± 13.1 nm radius), NUP107-GFP (from **b**, 219398 fitted localizations, 54.6 ± 11.9 nm radius), NUP96-SNAP (from **c**, 57297 fitted localizations, 55.9 ± 12.6 nm), NUP96-Halo (from **d**, 45143 fitted localizations, 56.2 ± 10.2 nm radius), NUP96-Anitbody (from **e**, 229069 fitted localizations, 43.5 ± 16 nm). See also **Supplementary Table 2.** Scale bars: 500 nm (overviews), 100 nm (individual NPCs and average).

Encouraged by the increased precision and reduced linkage error when targeting genetically-encoded probes with DNA-PAINT, we sought out to further optimize image acquisition conditions with respect to overall localization precision, sampling of single protein sites, and three-dimensional image acquisition. This allowed us to visualize single copies of NUP96 proteins (**Figure 3**) using the NUP96-Halo cell line, which we chose based on its superior performance in the 2D study presented above (smallest distribution width). An overview of a typical 3D DNA-PAINT dataset is shown in **Figure 3a**. Zooming in to some of the NPCs (**Fig. 3b**) reveals distinctive pairs of close-by “localization clouds” (arrows in **Fig. 3b**), which we attribute to single NUP96 proteins. To quantitatively asses the Euclidian distance of the two copies of NUP96 on the two cytoplasmic or nuclear rings of the NPC, we selected ~50 pairs in NPCs, aligned them on top of each other and subsequently carried out particle averaging with Picasso^4,25^. We then performed a crosssectional histogram analysis of the resulting sum image and fitted the distribution with two Gaussian functions (**Fig. 3c**). The fit yields a peak-to-peak distance of ~12 nm, well in agreement with the expected distance of NUP96 proteins on adjacent Y-complexes as derived from EM models^16^. Furthermore, each peak fit exhibits a standard deviation of only ~3 nm, highlighting the high localization precision and accuracy achievable with the combination of genetically-encoded tags with DNA-PAINT. Additionally, we measured the separation between the cytoplasmic and the nuclear rings for NUP96-Halo, yielding a distance of ~61 nm (**Fig. 3d**), which we could clearly resolve. The capability to separate the nuclear from the cytoplasmic side of the NPC is a necessity to indeed convince us, that the NUP pairs in each symmetry center (**Fig. 3b and c**) are indeed part of either the nuclear or cytoplasmic rings of the NPC. Furthermore, we obtained qualitatively and quantitatively similar results for the NUP96-SNAP cell line (**Supplementary Figure 7**).

**Figure 3.**
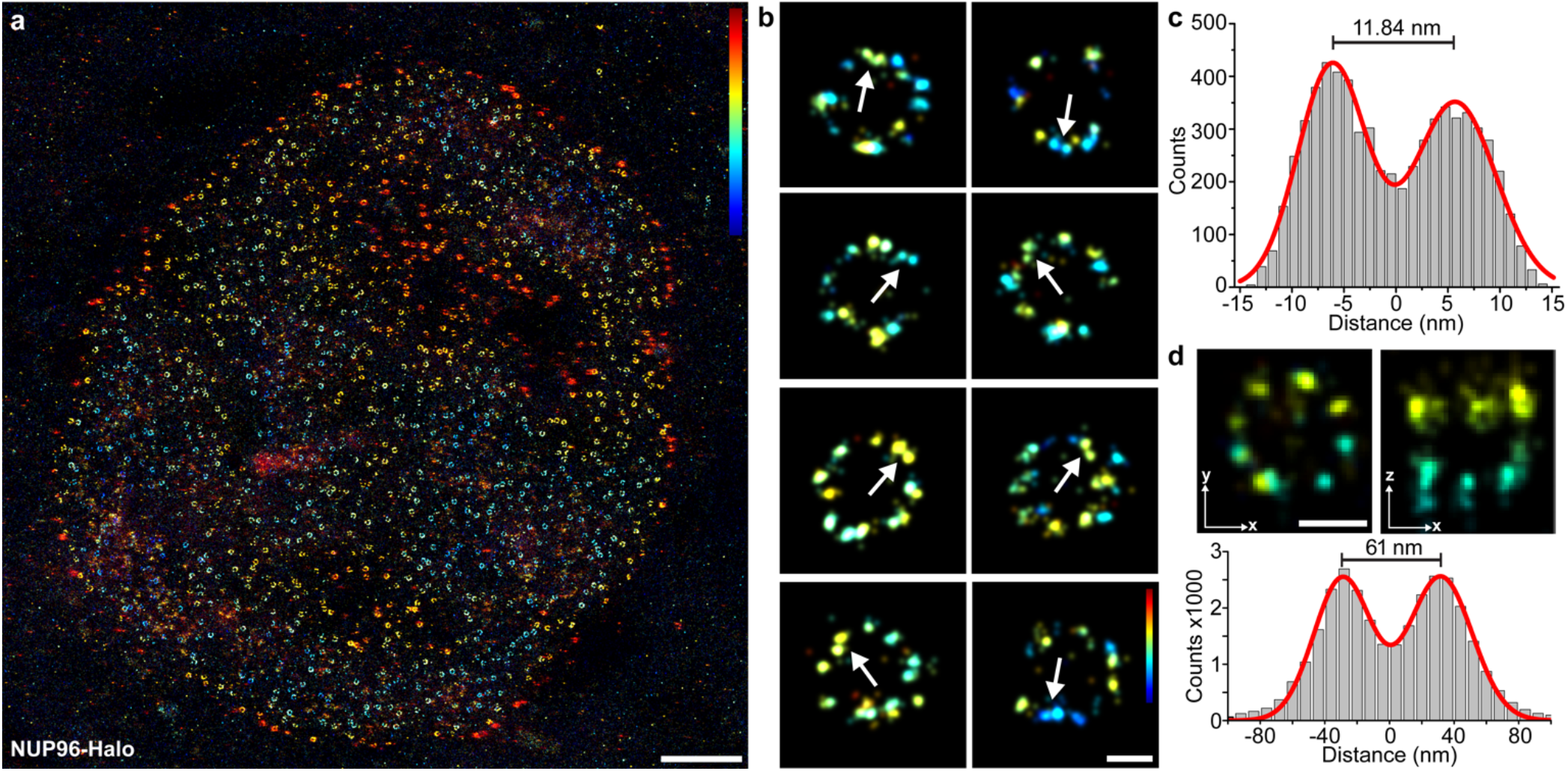
3D DNA-PAINT reveals single Halo-labeled NUP96 proteins in NPCs. (**a**) Overview image of NUP96-Halo imaged using 3D DNA-PAINT (color indicates height, range: −200 to 200 nm). (**b**) Selection of single NPCs. Arrows are highlighting two copies of NUP96 proteins in the same symmetry center of the same ring (i.e. at the same height) spaced ~12 nm apart from each other (color indicates height, range: −100 to 100 nm). (**c**) Cross-sectional histogram of 3D-averaged pairs (n = 45) of NUP96 proteins in single symmetry centers as highlighted by arrows in (**b**) yields ~12 nm distance between single proteins. (**d**) NUP96-Halo-labeled NPCs show the typical eightfold symmetry (xy-projection, left) and the organization in nuclear and cytoplasmic rings (xz-projection, right). Micrographs represent sum data from aligned NPCs (n = 31). **Bottom:** Crosssectional histogram of localizations in the xz-projection yields ~61 nm separation between cytoplasmic and nuclear rings. Scale bars: 2 μm (a), 50 nm (**b, d**).

In conclusion, we present a simple-to-implement, yet powerful approach to combine DNA-PAINT microscopy with genetically-encoded self-labeling tags. This brings DNA-based super-resolution microscopy to a level where it can provide a tool to investigate structural biology questions of higher order protein complexes in cells, thus far only amenable to EM. By keeping the advantages of fluorescence microscopy over EM, this now enables researchers to uncover single-molecule heterogeneities in complex protein assemblies on the level of single molecules. However, one of the main challenges in the field remains, which is a route to highly efficient labeling probes, ideally achieving >90 % labeling efficiency without requiring genetic engineering. Besides the availability of small scaffolds, like nanobodies^6^, affimers^26^, darpins^27^ or SOMAmers^28^ for binding the target of interest, novel approaches are necessary to tackle this challenge. These probes could include optimized host-guest systems^29^, direct transient binders^30^, or rationally-designed small proteins^31^. However, even with our currently achievable labeling efficiency, structural biology studies of multicomponent protein complexes in single cells using optical fluorescence microscopy are finally becoming a reality.

## Acknowledgements

We thank Bianca Sperl for experimental support. We thank Sandra Correia for assistance with generating the cell lines. This work has been supported in part by the German Research Foundation through the Emmy Noether Program (DFG JU 2957/1-1 to R.J.), the SFB1032 (project A11 to R.J.), the European Research Council through an ERC Starting Grant (MolMap; grant agreement number 680241 to R.J.) and an ERC Consolidator Grant (ERC CoG-724489 to J.R.), the Allen Distinguished Investigator Program through The Paul G. Allen Frontiers Group (to J.E. and R.J.), the National Institutes of Health Common Fund 4D Nucleome Program (Grant U01 EB021223 / U01 DA047728 to J.E. and J.R.), the European Molecular Biology Laboratory (EMBL; B.N., M.K., V.J.S., J.R., and J.E.) the Max Planck Foundation (to R.J.) and the Max Planck Society (to R.J.). T.S. and A.A. acknowledge support by the QBM graduate school. M.T.S. acknowledges support from the International Max Planck Research School for Molecular and Cellular Life Sciences (IMPRS-LS). V.S.J. acknowledges support by the Boehringer Ingelheim Fonds.

## Author contributions

T.S., M.T.S., F.S. and A.A. designed and performed experiments, analyzed data, and contributed to the writing of the manuscript. J.E., J.R., B.N., M.K. and V.J.S. created genetically-engineered cell lines. J.R. created GFP nanobodies. R.J. conceived and supervised the study, interpreted data, and wrote the manuscript. All authors reviewed and approved the manuscript. All requests for cell lines should be directed to Jan Ellenberg.

## Competing Interests

The authors declare no competing interests.

## ONLINE METHODS

### Tissue culture

McCoy’s 5A Medium modified (cat: 26600-023) was ordered from Gibco. Fetal Bovine Serum (FBS) (cat: 105000-64), 1× Phosphate Buffered Saline (PBS) pH 7.2 (cat: 20012-019), 0.05% Trypsin–EDTA (cat: 25300-054) and Penicillin-Streptomycin (cat: 15140-122) were purchased from Thermo Fisher Scientific. HeLa cells were purchased from the Leibniz Institute DSMZ (cat: ACC-57). Glass-bottomed 8-well μ-slides (cat: 80827) and sticky slide VI (cat: 80608) were obtained from ibidi. Falcon tissue culture flasks (cat: 734-0965) were ordered from VWR.

### Cell Fixation and immune staining

16% (w/v) Paraformaldehyde (cat: 28906) and DTT (cat: 20291) were purchased from Thermo Fisher Scientific. Triton X-100 (cat: 6683.1) and Ammonium chloride (cat: K298.1) was purchased from Roth. Bovine Serum Albumin (cat: A4503-10G) was ordered from Sigma-Aldrich. Halo- and SNAP-ligand-modified oligoes were custom-ordered from Biomers.net (see **Supplementary Table 3**). Primary monoclonal rabbit anti-NUP98 antibody (cat: 2598) was purchased from cell signaling. Primary monoclonal mouse anti-NUP98 was purchased from Santa Cruz Biotechnology (cat: sc-74578). Secondary polyclonal antibodies (cat: 711-005-152, 115-005-003) were purchased from Jackson ImmunoResearch.

### Cell imaging

EDTA 0.5 M pH 8.0 (cat: AM9261), Sodium Chloride 5 M (cat: AM9759) and Tris 1 M (cat: AM9856) were ordered from Ambion. Ultrapure water (cat: 10977-035) was purchased from Thermo Fisher Scientific. Potassium chloride (cat: 6781.1) was ordered from Roth. Sodium hydroxide (cat: 31627.290) was purchased from VWR. Glycerol (cat: G5516-500ML), Methanol (cat: 32213-2.5L), Protocatechuate 3,4-Dioxygenase Pseudomonas (PCD) (cat: P8279), 3,4-Dihydroxybenzoic acid (PCA) (cat: 37580-25G-F) and (+-)-6-Hydroxy-2,5,7,8-tetra-methylchromane-2-carboxylic acid (Trolox) (cat: 238813-5G) were purchased from Sigma-Aldrich. Dye modified DNA oligos were custom-ordered from MWG (see **Supplementary Table 4**). 90-nm-diameter Gold Nanoparticles (cat: G-90-100) were ordered from cytodiagnostics.

### Buffers

The imaging buffer was supplemented with: 100× Trolox: 100 mg Trolox, 430 μl 100 % Methanol, 345 μl 1M NaOH in 3.2 ml H_2_O. 40× PCA: 154 mg PCA, 10 ml water and NaOH were mixed and pH was adjusted 9.0. 100× PCD: 9.3 mg PCD, 13.3 ml of buffer (100 mM Tris-HCl pH 8, 50 mM KCl, 1 mM EDTA, 50 % Glycerol). Cell-imaging-buffer (buffer C): 1× PBS pH 8, 500 mM NaCl, 1× PCA, 1× PCD, 1× Trolox.

### PEG surface

PEG surfaces were prepared as previously reported^31^.

### GFP-Nanobody Labeling and Purification

GFP nanobodies were DNA-labeled as previously reported^32^. Nanobodies were concentrated via Amicon 10 kDa spin filters and buffer exchanged into 5 mM TCEP in 1× PBS + 3 mM EDTA at pH 7.5 mM TCEP in 1× PBS + 3 mM EDTA was then added to the GFP Nanobody and was incubated for 2 h at 4 °C on a shaker. Subsequently, Amicon 10 kDa Spin Filters were prewashed with 1× PBS, and Nanobody was buffer-exchanged into 1× PBS for 5× 5 min at 14 000×g and the volume was adjusted to 100 μl. DBCO-Maleimide Crosslinker was added in 20 molar excess in 5 μl to the GFP Nanobody and incubated overnight at 4 °C on a shaker. Crosslinker aliquots were stored at 40 mg/ml concentration in DMF. DBCO crosslinker was removed via 10 kDa Amicon Spin Filters for 5× 5min at 14 000× g. Azide-DNA was added to the GFP Nanobody crosslinker at 10× excess for 1 h at 20 °C. The final product was buffer exchanged into HisTrap binding buffer (20 mM Sodium Phosphate, 500 mM NaCl, 20 mM Imidazole) via Amicon 10 kDa Spin Filters. Purification from free DNA was performed using a GE Aekta purifier system and a HisTrap 1 ml column via a one-step purification scheme with 500 mM Imidazole and 500 mM NaCl. Peak fractions were afterwards concentrated and buffer-exchanged via Amicon 10 kDa spin filters.

### Antibody conjugation

Antibodies were conjugated to DNA-PAINT docking sites via maleimide-PEG2-succinimidyl ester chemistry as previously reported^33,34^ (see **Supplementary Table 3** for handle sequences).

### Cell culture

Hela cells and U2OS cells were passaged every other day and used between passage number 5 and 20. The cells were maintained in DMEM supplemented with 10 % Fetal Bovine Serum and 1 % Penicillin/Streptomycin. Passaging was performed using 1× PBS and Trypsin-EDTA 0.05 %. 24 h before immunostaining, cells were seeded on ibidi 8-well glass coverslips at 30,000 cells/well.

### Cell Fixation

Prefixation was performed with prewarmed 2.4 % Paraformaldehyde for 20 seconds followed by the permeabilization at 0.4 % Trion-X 100 for 10 seconds. Next, cells were fixed (main fixation) with 2.4 % for 30 min. After 3× rinsing with 1× PBS the cells were quenched with 50 mM Ammoniochloride (in 1× PBS) for 4 minutes. Then, cells were washed 3× with 1×PBS followed by incubation in 1× PBS for 5 minutes twice. Next, cells were stained with the corresponding ligand (see below). Finally, cells were washed 3× for 5 min in 1× PBS, incubated with 1:1 dilution of 90 nm gold particles in 1× PBS as drift markers, washed 3× 5 min and immediately imaged.

### Staining with SNAP

For SNAP-labeling, cells were incubated with 1 μM of SNAP-ligand-modified DNA oligomer in 0.5 % BSA and 1 mM DTT for 2 hours.

### Staining with HALO

For HALO-labeling, cells were incubated with 1 μM Halo-ligand-modified DNA oligomer in 3 % (w/v) BSA in 1× PBS for overnight at 4°C on a shaker.

### Staining with GFP

GFP-Nanobody staining was done in 3% BSA in 1xPBS at 4°C overnight on a shaker.

### Staining with Antibodies

Antibody staining was done in two steps. First, cells were incubated with primary antibody anti-NUP98 (1:100) in 3% BSA at 4°C PBS overnight. After three washes for 5 min with 1× PBS, the sample was incubated with the secondary antibody (dilution 1:100) at RT for 1 hours.

### Super-resolution microscope setup

Fluorescence imaging was carried out on an inverted Nikon Eclipse Ti microscope (Nikon Instruments) with the Perfect Focus System, applying an objective-type TIRF configuration with an oil-immersion objective (Apo SR TIRF 100×, NA 1.49, Oil). TIRF/Hilo angle was adjusted for highest signal to noise ratio when imaging. A 561 nm (200 mW, Coherent Sapphire) lasers was used for excitation. The laser beam was passed through cleanup filters (ZET561/10, Chroma Technology) and coupled into the microscope objective using a beam splitter (ZT561rdc, Chroma Technology). Fluorescence light was spectrally filtered with an emission filter (ET600/50m and ET575lp, Chroma Technology) and imaged on a sCMOS camera (Andor Zyla 4.2) without further magnification, resulting in an effective pixel size of 130 nm (sCMOS after 2×2 binning).

### Imaging conditions

**Figure 1c-e.** Imaging was carried out using an imager strand concentration of 300 pM (P3-Cy3B) in cell imaging buffer. 15,000 frames were acquired at 200 ms integration time. The readout bandwidth was set to 200 MHz. Laser power (@561 nm) was set to 30 mW (measured before the back focal plane (BFP) of the objective), corresponding to 0.7 kW/cm^2^ at the sample plane.

**Figure 1f-h.** Images were acquired with an imager strand concentration of 2 nM (P3-Cy3B imager) in cell imaging buffer. 40,000 frames were acquired at 200 ms integration time. The readout bandwidth was set to 200 MHz. Laser power (@561 nm) was set to 80 mW (measured at the back focal plane (BFP) of the objective). corresponding to 1.8 kW/cm^2^ at the sample plane.

**Figure 2a.** Images were acquired with an imager strand concentration of 2 nM of P3-Cy3B in in cell imaging buffer. 30,000 frames were acquired at 200 ms integration time and a readout bandwidth of 200 MHz. Laser power (@560 nm) was set to 50 mW (measured before the back focal plane (BFP) of the objective), corresponding to 1.1 kW/cm^2^ at the sample plane.

**Figure 2b.** Imaging was carried out using an imager strand concentration of 2 nM (P3-Cy3B) in cell imaging buffer. 30,000 frames were acquired at 200 ms integration time. The readout bandwidth was set to 200 MHz. Laser power (@561 nm) was set to 50 mW (measured before the back focal plane (BFP) of the objective), corresponding to 1.1 kW/cm^2^ at the sample plane.

**Figure 2c.** Images were acquired with an imager strand concentration of 2 nM (P3-Cy3B imager) in cell imaging buffer. 30,000 frames were acquired at 200 ms integration time. The readout bandwidth was set to 200 MHz. Laser power (@561 nm) was set to 50 mW (measured at the back focal plane (BFP) of the objective), corresponding to 1.1 kW/cm^2^ at the sample plane.

**Figure 2d.** Images were acquired with an imager strand concentration of 2 nM of P3-Cy3B in cell imaging buffer. 30,000 frames were acquired at 200 ms integration time. The readout bandwidth was set to 200 MHz. Laser power (@561 nm) was set to 50 mW (measured at the back focal plane (BFP) of the objective), corresponding to 1.1 kW/cm^2^ at the sample plane.

**Figure 2e.** Imaging was carried out using an imager strand concentration of 300 pM (P3-Cy3B) in cell imaging buffer. 30,000 frames were acquired at 200 ms integration time. The readout bandwidth was set to 200 MHz. Laser power (@561 nm) was set to 50 mW (measured before the back focal plane (BFP) of the objective), corresponding to 1.1 kW/cm^2^ at the sample plane.

**Figure 3a, b.** Images were acquired with an imager strand concentration of 2 nM (P3-Cy3B imager) in cell imaging buffer. 100,000 frames were acquired at 200 ms integration time. The readout bandwidth was set to 200 MHz. Laser power (@561 nm) was set to 40 mW (measured at the back focal plane (BFP) of the objective), corresponding to 1.0 kW/cm^2^ at the sample plane.

For all imager strand sequences see **Supplementary Table 4.**

### 3D DNA-PAINT calibration using latex microspheres

The 3D look-up table was measured as previously reported^35^. In short, first an ibidi sticky slide VI was assembled with the pegylated coverslip. Then, 50 μl of 1:10 avidin coated microspheres diluted in 1× PBS were flown into the ibidi sticky slide chamber with the prepared PEG-Biotin surface and incubated for 10 min. Then the chamber was washed using 180 μl of 1x PBS. Second, 500 nM biotinylated oligonucleotides (10 nt, P1 docking site sequence, **Supplementary Table 4**) was then flown into the chamber and incubated for 10 min. Next, the chamber was washed with 180 μl of 1× PBS. Next, the chamber was incubated with 1:10 dilution of 90 nm gold particles in 1× PBS as drift markers for 5 min and subsequently washed with 80 μl 1× PBS. Finally, 180 μl imaging buffer with dye-labeled imager strands was flown into the chamber. 500 pM Cy3B labeled imager with sequence P1 and 1× PCA, 1× PCD, 1× Trolox in buffer C was used. Latex microspheres attached to the PEG surface were identified using bright-field illumination and the radius was measured. The recoded latex microsphere data using DNA-PAINT was reconstructed using two-dimensional gaussian fitting. Lateral drift correction was performed using the gold nanoparticle. Gaussian width (sigma x and sigma y) were averaged in radial sections and linked to the corresponding z height to gather the calibration data^36^. Finally, the calibration data was fitted using sixth degree polynomial fit to generate the look-up table.

### Image analysis

Raw fluorescence data was subjected to spot-finding and subsequent super-resolution reconstruction using the ‘Picasso’ software package^34^. x, y and z Drift correction was performed with a redundant cross-correlation and gold particles as fiducials.

### Radius analysis

To determine the radius of NPCs, picked NPCs were averaged using the ‘Picasso:average3’ module as previously described^34^. In brief, localizations of particles are aligned on top of each other by rendering them and using cross-correlation to determine displacement. To account for ring-like structures, a 100× symmetry was set. Each dataset was averaged with the following oversampling settings: 3× 15, 1× 20, 1× 40. Based on the resulting “superparticle”, the center of mass was determined.

The localizations were subsequently transformed into polar coordinates with the center of mass as the center point. The radius was calculated by taking the median of the polar coordinate distances.

