## Supplementary Information for "Direct visualization of single nuclear pore complex proteins using genetically-encoded probes for DNA-PAINT"


| | |
|---|---|
| **Supplementary Figure 1** | **Overview of NUP107-SNAP, n=398** |
| **Supplementary Figure 2** | **Overview of NUP107-GFP, n=486** |
| **Supplementary Figure 3** | **Overview of NUP96-SNAP, n=288** |
| **Supplementary Figure 4** | **Overview of NUP96-Halo, n=191** |
| **Supplementary Figure 5** | **Overview of NUP96-AB, n=185** |
| **Supplementary Figure 6** | **NUP96-AB and NUP107-NB** |
| **Supplementary Figure 7** | **NUP96-SNAP 3D DNA-PAINT** |
| **Supplementary Table 1** | **Imaging Parameters** |
| **Supplementary Table 2** | **Imager Sequences** |
| **Supplementary Table 3** | **Handle Sequences** |
| **Supplementary Table 4** | **NPC radius quantification** |
| **Supplementary References** | |

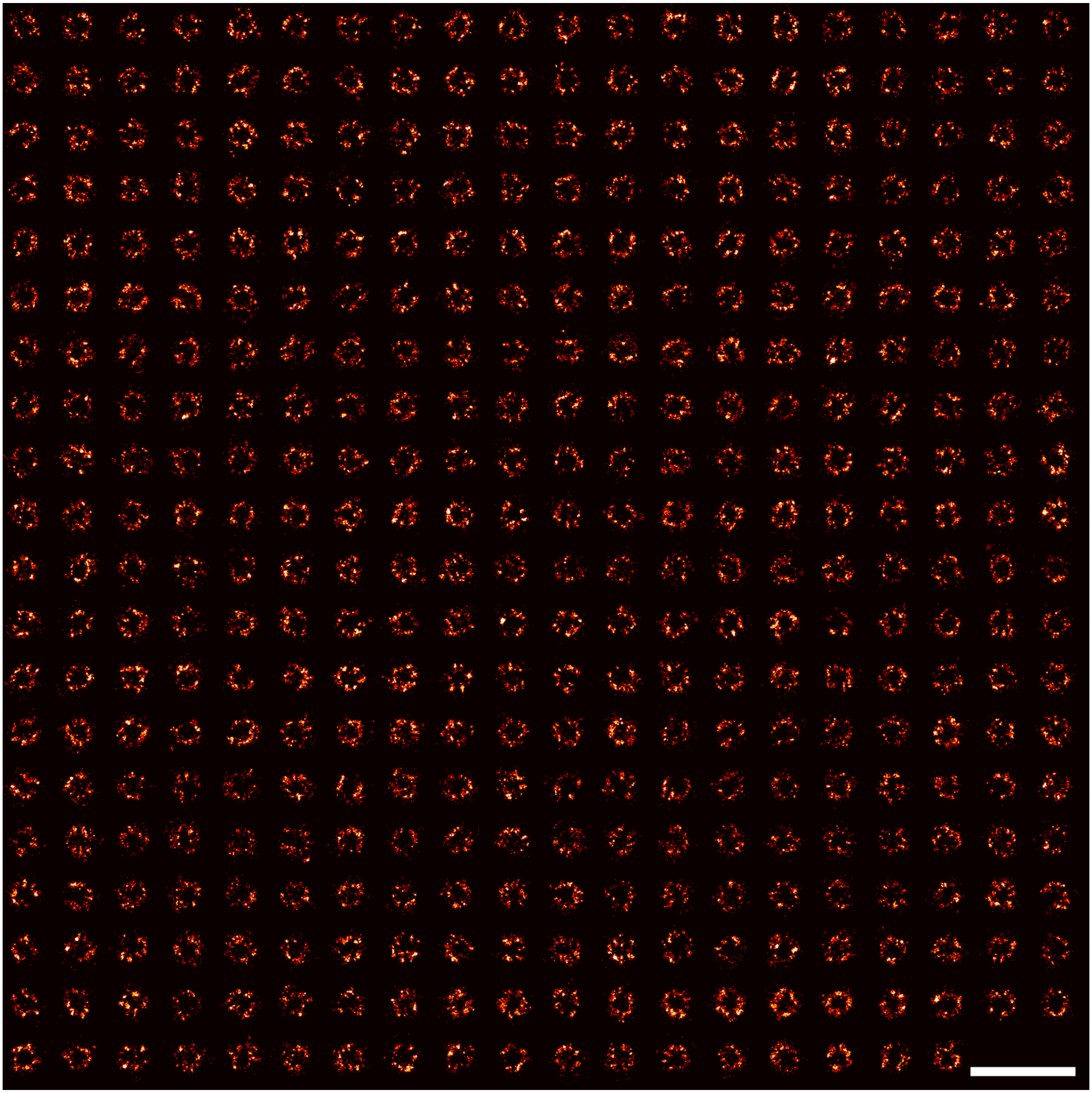

**Supplementary Figure 1 | Overview of NUP107-SNAP, n=398.** Scale bar: 500 nm.

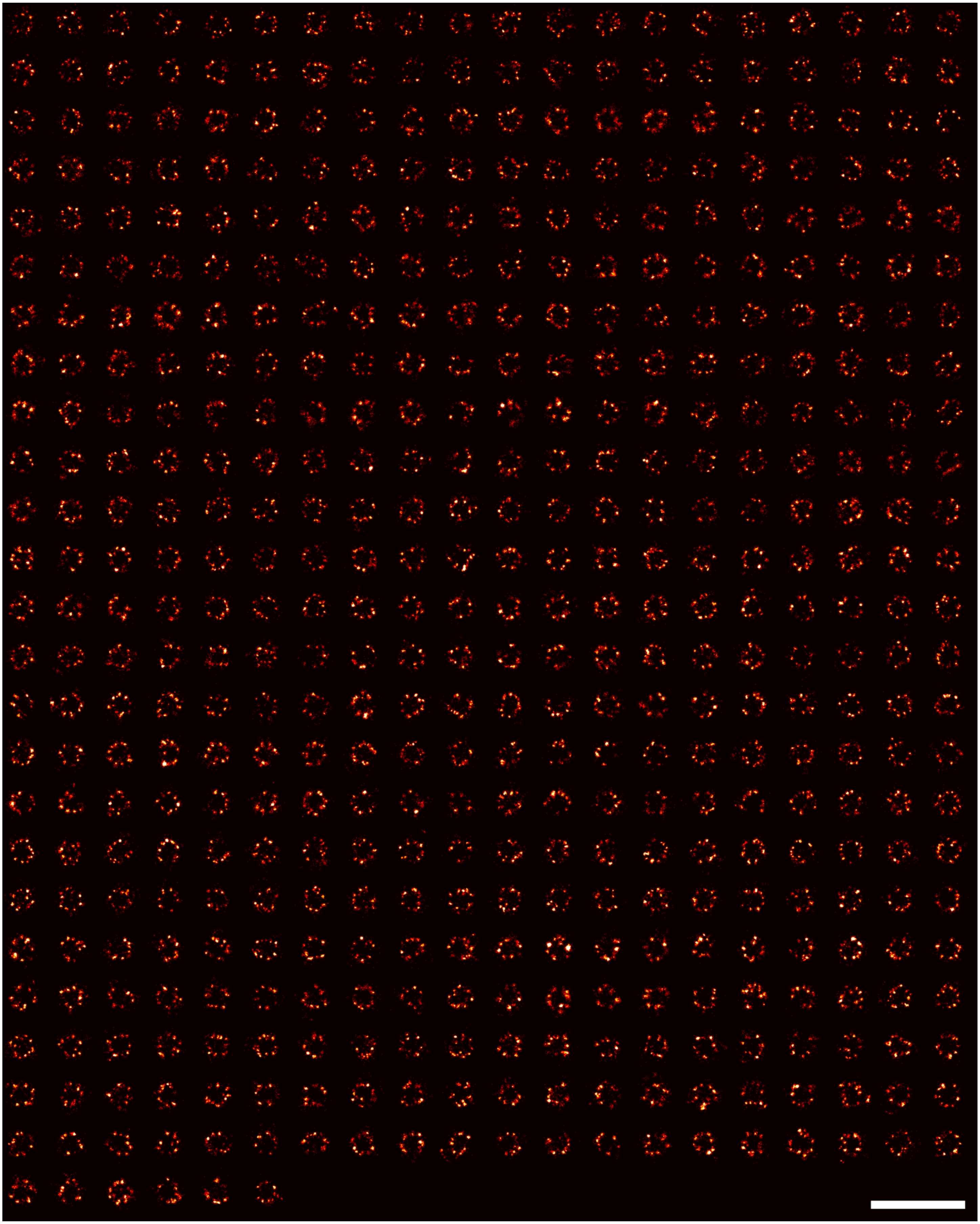

**Supplementary Figure 2 | Overview of NUP107-GFP, n=486.** Scale bar: 500 nm.

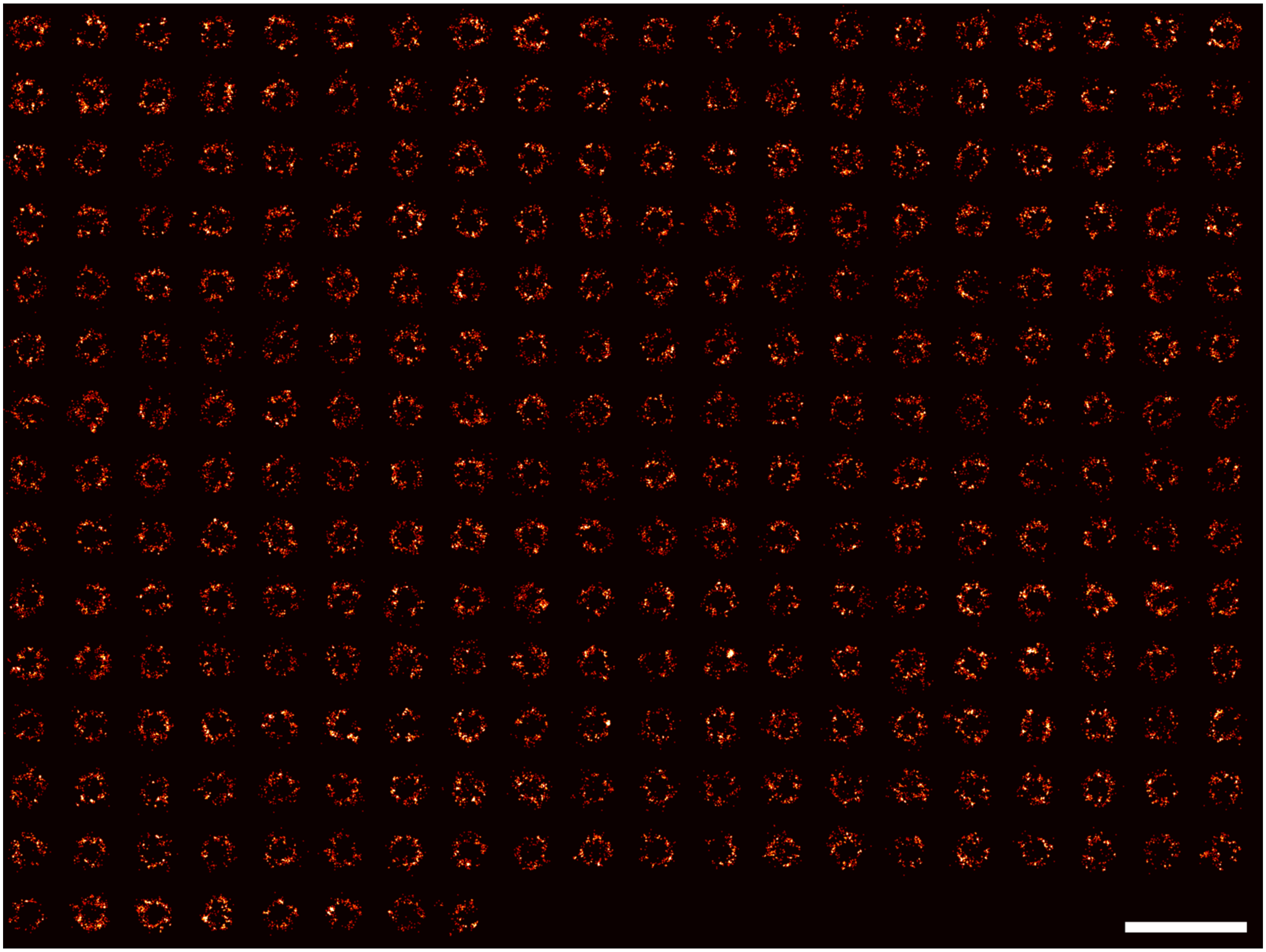

**Supplementary Figure 3 | Overview of NUP96-SNAP, n=288.** Scale bar: 500 nm.

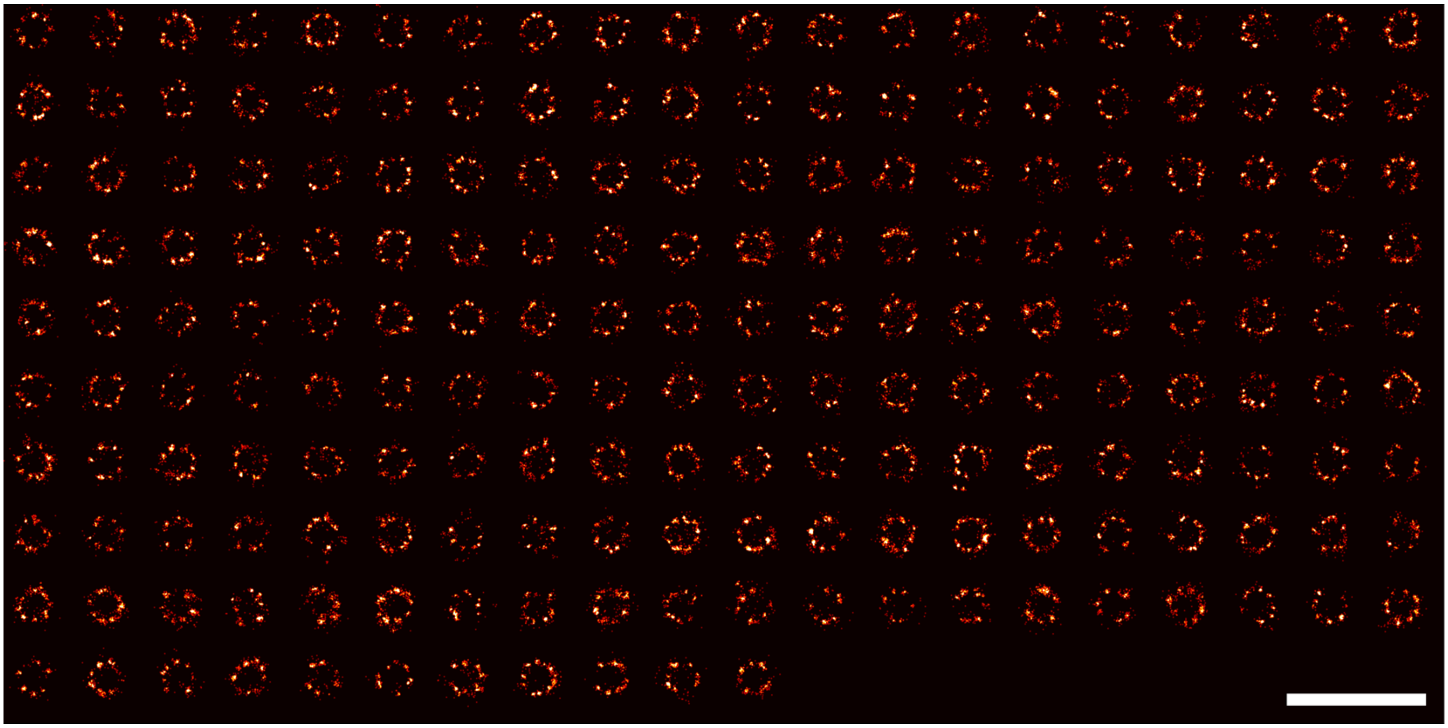

**Supplementary Figure 4 | Overview of NUP96-Halo, n=191.** Scale bar: 500 nm.

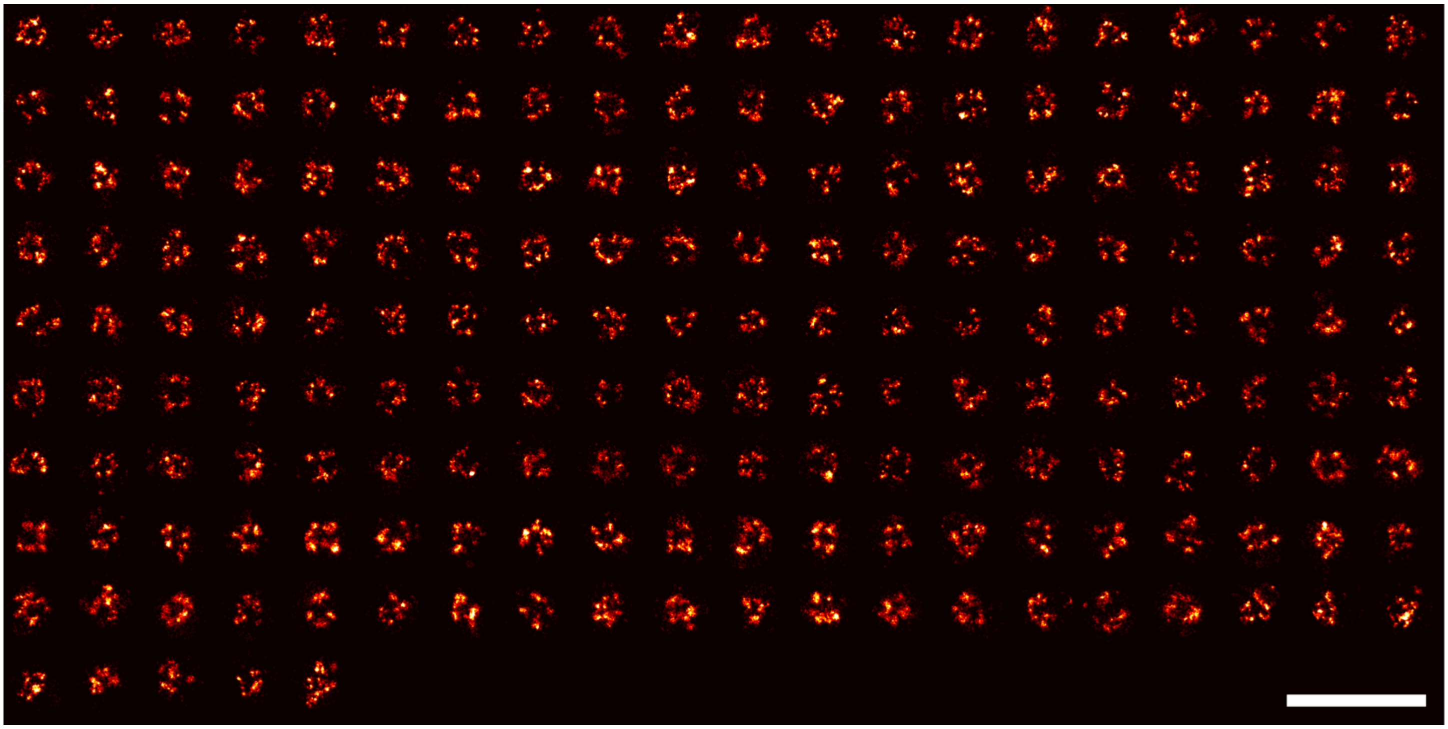

**Supplementary Figure 5 | Overview of NUP96-AB, n=185.** Scale bar: 500 nm.

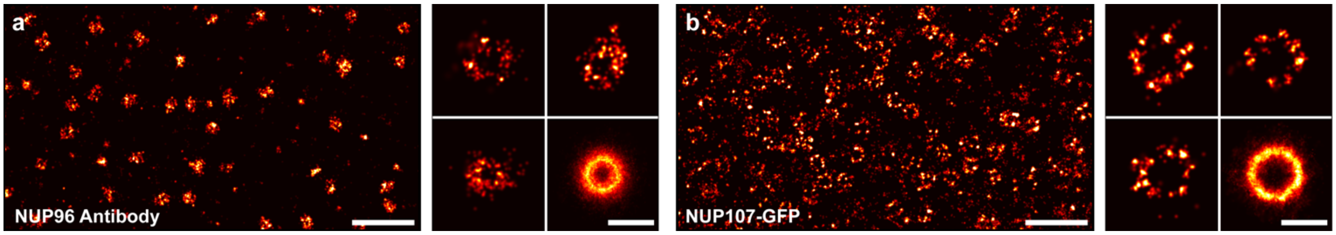


**Supplementary Figure 6 | NUP96-AB and NUP107-NB.** Scale bars: 500 nm (overviews), 100 nm (individual NPCs and average).

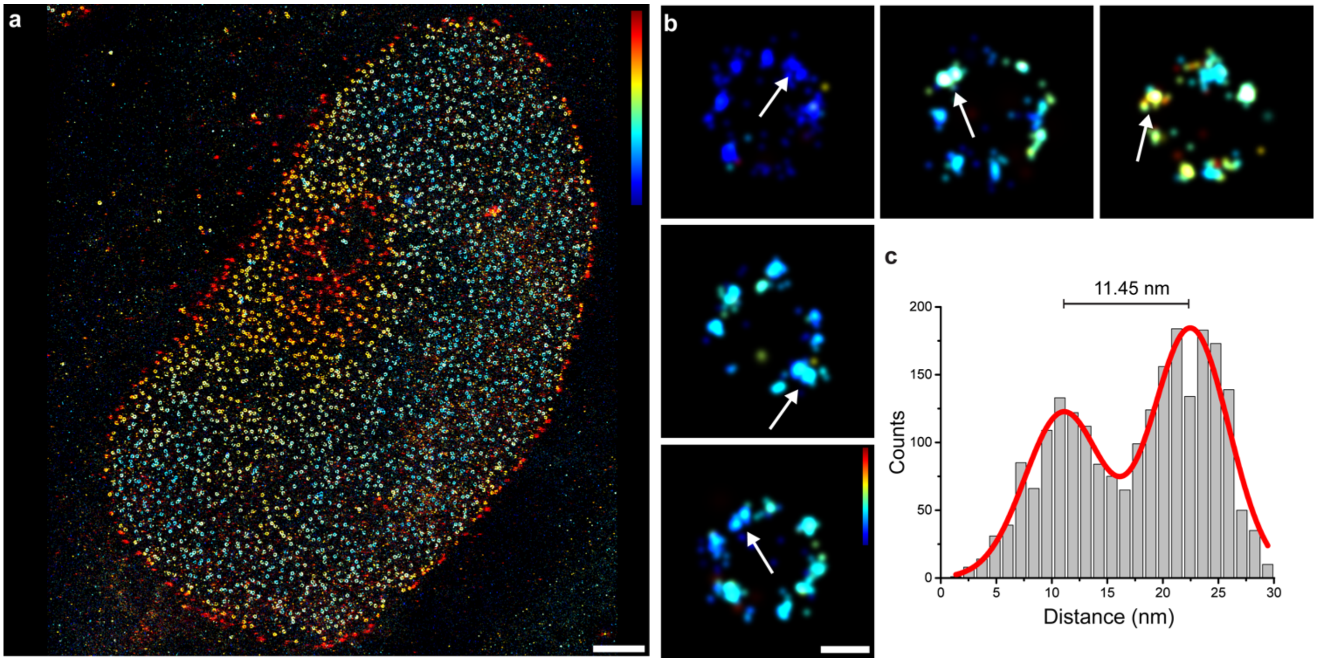


**Supplementary Figure 7 | NUP96-SNAP 3D DNA-PAINT.** (**a**) 3D DNA-PAINT overview image of NPCs labeled via Nup96-SNAP (color indicates height, range: -200 to 200 nm). (**b**) Selection of single NPCs. Arrows are highlighting two copies of NUP96 proteins in the same symmetry center of the same ring (i.e. at the same height) spaced ~12 nm apart from each other (color indicates height, range: -100 to 100 nm). (**c**) Cross sectional histogram of 3D-averaged pairs (N = 27) of NUP96-SNAP proteins in single symmetry centers as highlighted in (**b**). Scale bars: 2 µm (**a**), 50 nm (**b**).

Imaging was carried out using an imager strand concentration of 2 nM (P3-Cy3B). 50k frames were acquired at 200 ms integration time. The readout bandwidth was set to 200 MHz. Laser power (@561 nm) was set to 40 mW (measured before the back focal plane (BFP) of the objective). This corresponds to 1 kW/cm$^2$ at the sample plane.

**Supplementary Table 1 | Imaging parameters**

| Dataset | Parameters | Power @561 nm | NeNA precision |
|---|---|---|---|
| Figure 1c,d,e | 300ms, 2D, 15k Frames, 5nM. P3* | 0.7 kW/cm$^2$ | 6.3 nm |
| Figure 1f,g,h | 200ms, 2D, 30k Frames, 2nM, P3* | 1.8 kW/cm$^2$ | 6.9 nm |
| Figure 2a and SI Figure 1 | 200 ms, 2D, 30k Frames, 1 nM, P3* | 1.1 kW/cm$^2$ | 3.6 nm |
| Figure 2b and SI Figure 2 | 200 ms, 2D, 30k Frames, 2nM, P3* | 0.6 kW/cm$^2$ | 6.0 nm |
| Figure 2c SI Figure 3 | 200 ms, 2D, 30k Frames, 1 nM, P3* | 1.1 kW/cm$^2$ | 2.9 nm |
| Figure 2d and SI Figure 4 | 200 ms, 2D, 30k Frames, 1 nM, P3* | 1.1 kW/cm$^2$ | 4.5 nm |
| Figure 2e and SI Figure 5 | 200ms, 2D, 30k Frames, 2 nM, P3* | 0.6 kW/cm$^2$ | 4.5 nm |
| Figure 3 | 200ms, 3D, 100k Frames, 2nM, P3* | 1 kW/cm$^2$ | 6.0 nm |
| SI Figure 6a | 200 ms, 2D, 30k Frames, 300 pM, P3* | 1.1 kW/cm$^2$ | 2.2 nm |
| SI Figure 6b | 200 ms, 2D, 30k Frames, 1 nM, P3* | 1.1 kW/cm$^2$ | 6.0 nm |
| SI Figure 7 | 200 ms, 3D, 50k Frames, 2 nM, P3* | 1 kW/cm$^2$ | 5.1 nm |

**Supplementary Table 2 | Imager sequences**

| Imager name | Sequence | 5'-mod | 3'-mod | Vendor |
|---|---|---|---|---|
| P1* | CTAGATGTAT | None | Cy3b | Eurofins Genomics |
| P3* | GTAATGAAGA | None | Cy3b | Eurofins Genomics |

**Supplementary Table 3 | Handle sequences**

| Handle Name | Sequence | 5'-mod | 3'-mod | Vendor |
|---|---|---|---|---|
| P1 | TTATACATCTA | BG (Snap Ligand) | None | Biomers.net |
| P3 | TTTCTTCATTA | BG (Snap Ligand) | None | Biomers.net |
| P1 | TTATACATCTA | Halo Ligand | None | Biomers.net |
| P3 | TTTCTTCATTA | Halo Ligand | None | Biomers.net |
| P1 | TTATACATCTA | Thiol (for AB conjugation) | None | Eurofins Genomics |
| P3 | TTTCTTCATTA | Thiol (for AB conjugation) | None | Eurofins Genomics |
| P1 | TTATACATCTA | Biotin | None | Eurofins Genomics |

**Supplementary Table 4 | NPC radius quantification**

| Dataset | Median (nm) | Mean (nm) | Std (nm) | # Pores | # Locs |
|---|---|---|---|---|---|
| SNAP Nup107 | 53.7 | 54.2 | 13.1 | 398 | 127773 |
| GFP-NB Nup107 | 54.6 | 54.8 | 11.9 | 486 | 219398 |
| SNAP Nup96 | 55.9 | 56.5 | 12.6 | 288 | 57297 |
| HALO Nup96 | 56.2 | 56.6 | 10.2 | 191 | 45143 |
| AB Nup98 | 43.5 | 45.2 | 16 | 185 | 229069 |
